# Multi-scale predictions distinctively modulate tone perception in schizophrenia patients with auditory verbal hallucinations

**DOI:** 10.1101/2020.07.15.204495

**Authors:** Fuyin Yang, Hao Zhu, Lingfang Yu, Weihong Lu, Chen Zhang, Xing Tian

## Abstract

Auditory verbal hallucinations (AVHs) are one of the most pronounced symptoms that manifest the underlying mechanisms of deficits in schizophrenia. Cognitive models postulate that malfunctioned source monitoring incorrectly weights the top-down prediction and bottom-up sensory processing and causes hallucinations. Here, we investigate the featural-temporal characteristics of source monitoring in AVHs. Schizophrenia patients with and without AVHs, and healthy controls identified target tones in noise at the end of tone sequences. Predictions of different timescales were manipulated by either an alternating pattern in the preceding tone sequences, or a repetition between the target tone and the tone immediately before. The sensitivity index, d’, was obtained to assess the modulation of predictions on tone identification. We found that patients with AVHs showed higher d’ when the target tones conformed to the long-term regularity of alternating pattern in the preceding tone sequence than that when the targets were inconsistent with the pattern. Whereas, the short-term regularity of repetitions modulated the tone identification in patients without AVHs. Predictions did not influence tone identification in healthy controls. These findings suggest that malfunctioned source monitoring in AVHs heavily weights predictions to form incorrect perception. The weighting function in source monitoring can extend to the process of basic tonal features, and predictions at multiple timescales differentially modulate perception in different clinical populations. These collaboratively reveal the featural and temporal characteristics of weighting function in source monitoring of AVHs and suggest that the malfunctioned interaction between top-down and bottom-up processes might underlie the development of auditory hallucinations.

**Highlights:** Malfunctioned source monitoring incorrectly weights the top-down prediction and bottom-up sensory processing underlie pathogenesis of auditory verbal hallucinations in schizophrenia.

The weighting function in top-down predictions and bottom-up sensory processing can extend to tonal features.

Predictions at multiple timescales differentially modulate perception in different clinical schizophrenia populations.

## Introduction

Auditory verbal hallucinations (AVHs) are symptoms of hearing voices in the absence of external stimuli (Stephane *et al.*, 2001). Around 50%-70% of people who are diagnosed with schizophrenia experience AVHs (Meltzer, 1992, Waters, 2012). Antipsychotic medication such as olanzapine, risperidone and quetiapine that block D2-receptors in the brain are the most effective drugs available to treat AVHs (Farde, 1997, Sommer *et al.*, 2012). However, weight gain and sedation are serious side effects associated with antipsychotic medications (Leucht *et al.*, 2013). Medications may expose patients to metabolic complications and result in treatment non-adherence. Moreover, about 25% patients are resistant to standard antipsychotic treatment (Shergill *et al.*, 1998). Noninvasive neuro-stimulation techniques have been tested as a new treatment option for AVHs (Hoffman *et al.*, 2000). Repetitive transcranial magnetic stimulation (rTMS) and transcranial direct current stimulation (tDCS) are two noninvasive techniques that are recently introduced to treat AVHs (Mondino *et al.*, 2015, Priori *et al.*, 2009). The rTMS and tDCS, as promising treatment options (Brunelin *et al.*, 2012, Sommer *et al.*, 2012), show a moderate effect size in reduction of AVHs frequency (Brunoni *et al.*, 2011, Yang *et al.*, 2019). That the treatment effects depend on the stimulation protocols and cortical targets (Kindler *et al.*, 2013, Yang *et al.*, 2019) highlights the necessity of understanding AVHs from a cognitive neuroscience perspective.

The cognitive models postulate that AVHs may result from a process in which inner or sub-vocal speech is misidentified as externally caused (Frith and Done, 1989). Such source monitoring account of AVHs requires an internally generated source. Prediction has been proposed as an algorithm that induce this internal source (Moseley *et al.*, 2013). Prediction refers to the set of processes that are based on the information of memory, knowledge, and belief to generate representations of future events (Moulton and Kosslyn, 2009, Schacter *et al.*, 2012, Szpunar *et al.*, 2014, Teufel and Fletcher, 2020). For example, similar representations as perception can be constructed based on retrieving memory of past experiences and regularities without external stimulations (Tian and Poeppel, 2014, Tian *et al.*, 2016, Wheeler *et al.*, 2000). These internally constructed representations from prediction could be the critical source in the cognitive monitoring account of AVHs.

Combining the internal source of prediction and the external source of sensory analysis, the computation of source monitoring can be quantified as a weighting function between prediction and perception. Numerous studies have suggested that the top-down prediction interacts with bottom up sensory processes to shape perception (Bentall and Slade, 1985, Corlett *et al.*, 2019, Ma and Tian, 2019). Predictions can economically balancing the cognitive resources for processing the adapted perception and detecting the unexpected events (Tian and Huber, 2013). Breakdown in predictive function means individuals are less likely to attend to effective indicators of upcoming sensation (Keefe and Kraus, 2009). The unbalanced weighting between sources from prediction and sensory input could cause AVHs (Aleman *et al.*, 2003). For example, participants with severe hallucinations significantly increase gain over prior predictions in ambiguous perceptual situations (Cassidy *et al.*, 2018), suggesting that a relatively higher priority is assigned to top-down factors in determining the final percepts (Behrendt, 1998). To an extreme, abnormal top-down prediction processes in patients overwhelm the auditory input (Aleman *et al.*, 2003, Dima *et al.*, 2010). The malfunctioned weighting in the source monitoring is consistent with the framework of Bayesian inferences, share the assumption that AVHs are induced when sensory predictions are activated without sensory input, or these predictions are not properly deactivated and incorrect replace the sensory analysis (Powers *et al.*, 2017).

What perceptual features and temporal characteristics the weighting function are operating on are two crucial aspects to understand the source monitory of AVHs. First, source monitoring weights and balances multiple levels of features across different sources to establish coherent percepts. Previous studies have demonstrated the influences of top-down prediction at the semantic and phonological levels in healthy subjects (Davis and Johnsrude, 2007). Recently, the effects of top-down prediction on sensory analysis have extended to lower and basic sound attributes, such as pitch and loudness (Tian *et al.*, 2018, Tian and Poeppel, 2014). However, whether the weighting function in source monitoring of AVHs can extend to basic level attributes is still in debate. Some studies found that the severity of hallucination-prone was correlated with errors that were induced by semantic priming but not with phonological priming (Vercammen *et al.*, 2008). Other studies provided preliminary evidence of increased top-down influences for tonal stimuli (Aleman *et al.*, 2003). In this study, we investigated whether AVHs patients increased top-down influences in the processing of tones. That is, we aim to answer whether the source monitoring in AVHs only incorrectly weights the higher level features or have a ubiquitous weighting function that applies on sound attributes of all levels.

Second, prior information is available at multiple timescales and facilitate information processing across time (Fuster, 1997). For example, speech processing may operate at two distinct timescales (Giraud and Poeppel, 2012, Teng *et al.*, 2017). Multiple levels of prior information could help comprehension of linguistic information ranging from phonemes, to words, to sentences, and to paragraphs (Ding *et al.*, 2016, Hickok and Poeppel, 2007, Teng *et al.*, 2020). Memory of recent events integrates information over milliseconds, seconds, and minutes to form predictions at multiple timescales that continuously support the processing of incoming information (Hasson *et al.*, 2015). Would the source monitoring of AVHs weight predictions at multiple timescales differently?

Together, this study investigated the featural and temporal characteristics of the weighting function in source monitoring of AVHs. Specifically, we hypothesized that AVHs are caused by an incorrect weighting of top-down predictions, distorting the balance between bottom-up and top-down processes. This distorted balance of weighting sources in AVHs may influence the processing of basic speech features such as tones, and predictions at different timescales may modulate the bias differently.

In this study, we manipulated the long-term regularity and immediate repetitions in sequences of tones to investigate the featural and temporal characteristics of the weighting function in source monitoring of AVHs. Sequences of tones were presented either in an alternating pattern (long-term regularity) or randomly with the possibility that the last tone repeated the immediate preceding one (immediate repetitions). Three groups of participants, patients with and without AVHs as well as healthy controls, were asked to identify the last tone that was embedded in noise. Perceptual sensitivity, d’, was obtained based on the signal detection theory (SDT). According to our hypothesis that patients with AVHs may confuse the sources of their memory-based predictions and the sensory processing of tonal features, we predicted that the d’ of tone identification in the group of AVHs would be modulated by the manipulations in the preceding tone sequences. Moreover, the modulation effects would be different for the long-term and short-term predictions across groups.

## Methods

### Participants

Thirty-two (14 males) patients, who matched a DSM-V diagnosis of schizophrenia and were currently experiencing AVHs (AVHs patients) without concomitant hallucinations in other modalities, were recruited from Shanghai Mental Health Center. Furthermore, twenty-nine (12 males) patients met DSM-V diagnosis of schizophrenia who had never experienced AVHs (non-AVHs patients) were recruited from the same hospital. Two experienced psychiatrist independently diagnosed each patient, and the diagnosis was confirmed by the Structured Clinical Interview for DSM-V (SCID). All patients were receiving atypical antipsychotic medications and were clinically stable.

Thirty healthy Chinese subjects (9 males) were recruited as the control group from the local communities and schools in Shanghai. A clinical psychiatrist assessed these healthy subjects’ current mental status and any personal and family history of mental disorders. Moreover, any subject with potential psychiatric morbidity was excluded from the control group after the psychiatrist’s unstructured interviews. None of the healthy subjects had any family history of psychiatric disorders or physical diseases.

All participants were in the age range of 18-45 years old, right-handed, and without any substance abuse records.

The study was approved by the Institutional Review Board of the New York University Shanghai and the Institutional Ethics Committee at Shanghai Mental Health Center. All participants provided signed, informed consent before their participation in the study.

### Clinical measures

Demographic data were collected from patients and healthy controls. Four psychiatrists, who were blind to the study, assessed the patient’s psychopathology using the Positive and Negative Syndrome Scale (PANSS). The PANSS measures both the presence and severity of Positive, Negative and General symptoms on a 7-point scale. AVHs severity was rated from the P3 of the PANSS scale, with higher ratings indicating an increase in AVHs severity. Non-AVHs patients had a rating of 1 in the P3 factor score, indicating that the symptom was absent. Additionally, AVHs level of current AVHs patients were assessed using the 7-item Auditory Hallucinations Rating Scale-AHRS (Hoffman *et al.*, 2003). To ensure consistency and reliability of PANSS and AHRS, paired ratings between two psychiatrists for the same patient assessment were compared at each of the repeated assessments. All paired rating had a correlation coefficient >0.8 on the PANSS and AHRS total scores.

### Experimental design

#### Materials

Two different Mandarin tones of vowels /a/ (/ā/ and /á/) were synthesized via the NeoSpeech engine (NeoSpeech, 2012) with a female voice. Both Mandarin tones were 377ms in duration and scaled to 75dB SPL in intensity using Praat software.(Boersma, 2002) Additional two stimuli were created by adding white noise to the two mandarin tones. The signal-to-noise ratio was determined at individual level during a pre-test. All stimuli were digitized at 44.1kHz sampling rate and 16bit bitrate. These auditory stimuli were delivered through Sennheiser HD 280 Pro headphones. The volume was adjusted to a comfortable level for each participant and kept consistent throughout experiment for all stimuli.

#### Threshold Test Procedures

This procedure was composed of two different steps. First, participants participated in a pre-test that measured a threshold for the detection of two Mandarin tones (/ā/ or /á/) in white noise. This pre-test session consisted of 200 trials. At the beginning of the trial, a visual cue was presented for 500ms. After the offset of the visual cue, one of the auditory stimuli in noise was presented. White noise was 1s in duration. The auditory stimulus was presented 100 ms after the onset of the white noise and lasted for 377 ms. The intensity of the white noise changed trial by trial given by the Bayesian adaptive “PSI” staircase method (Kontsevich and Tyler, 1999) while the intensity of the vowels fixed at 75dB. The Psi-staircase assumed a log-Weibull (Gumbel) function with a non-zero (2%) attentional lapse rate (Lambda) and a 5% guess rate (Gamma). Two randomly interleaved Psi-staircases for the two auditory stimuli were created with 100 trials per staircase. Participants were required to provide a perceptual judgment of the tone in a two-alternative farced-choice (2AFC) paradigm. Participants took a break of a few minutes after every 50 trials. The threshold for each tone was determined by the 75% accuracy point in the fitted the psychometric curves for each participant. The signal-to-noise ratio (SNR) used in the main experiment was determined by the (fixed) intensity of signals to the threshold intensity of noise.

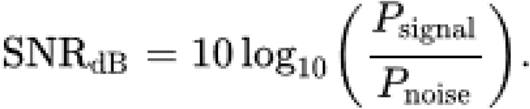

Last, we ran a confirmation test. Participants judged twenty trials (ten for each of tone) with the SNR determined in the pre-test. When the number of correctly judgments was seven or eight in each of tone, Participants were considered pass the test. Only participants who passed the confirmation test could proceed to the next procedure.

#### Main Procedure

After determining an appropriate threshold of each tone, participants proceeded to the main experiment in which they heard a sequence of tones and make judgement to the last tone in noise.

At the beginning of a trial, a fixation appeared in the center of the screen for 500 msec. After the onset of the visual cue, participants passively heard four to seven clean /ā/ or /á/ Mandarin tone in a sequence. The duration of each tone was 377ms. The inter-stimulus interval (ISI) was 623ms. Therefore, the stimulus onset asynchrony (SOA) was 1s. Trails with different number of tones were randomly presented. The last stimulus in a trial was always a tone in noise with the individual measured SNR in the pre-test. The target was randomly selected from /ā/ or /á/, and was presented in noise in the same way as in the pre-test 100 ms after the onset of 1s-long white noise and lasted for 377ms. Participants judged whether the tone in the noise was /ā/ or /á/ by pressing one of two buttons (Figure 1). The probability of each tone in the first clean tone position, in the last clean tone position, and in the noise was same.

**Figure 1.**
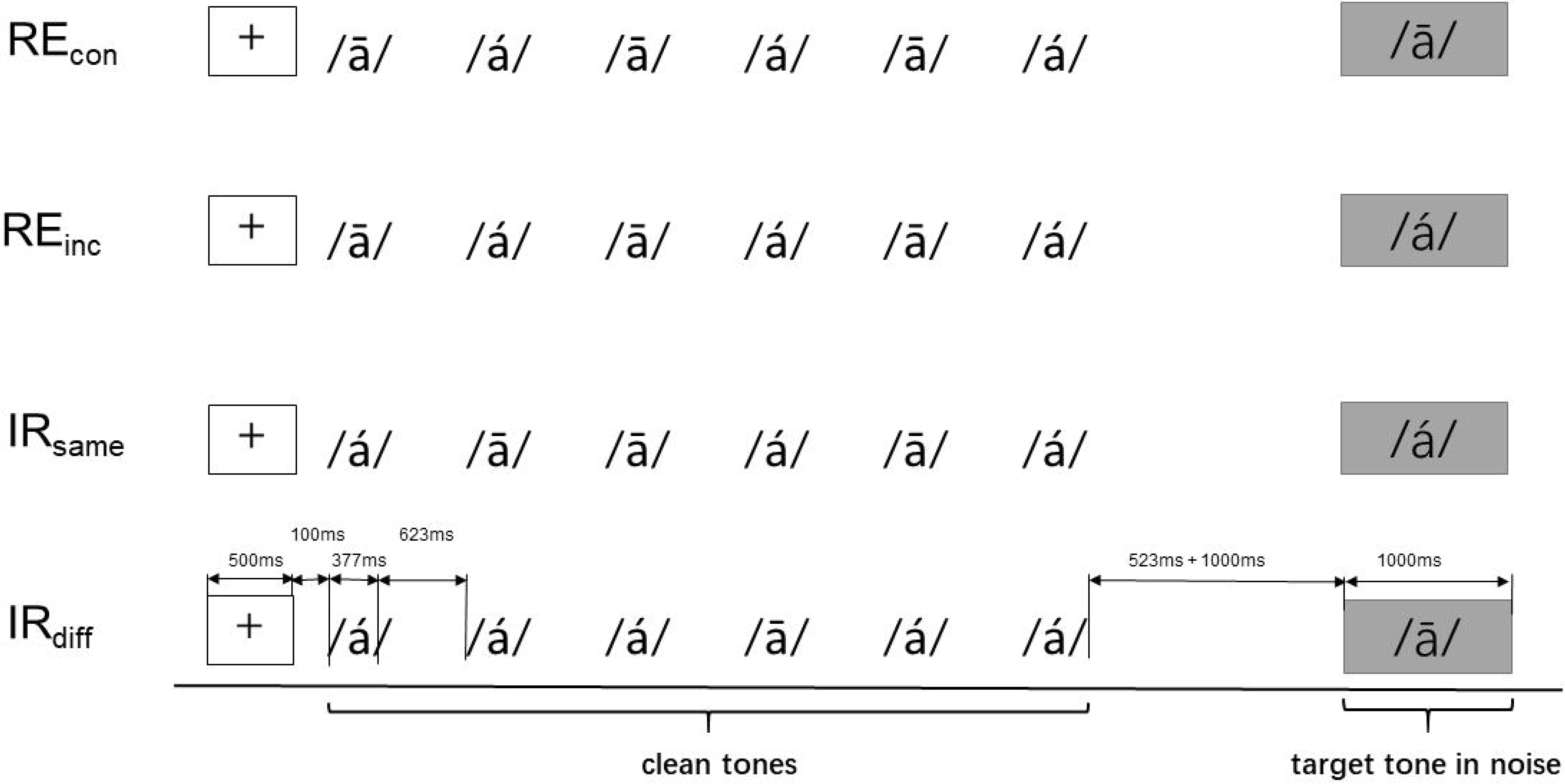
Schematic description of experimental procedures. At the beginning of a trial, a fixation appeared in the center of the screen for 500 msec. After the offset of the visual cue, participants passively heard four to seven clean /ā/ or /á/ Mandarin tones in a sequence. The duration of each tone was 377ms. The inter-stimulus interval (ISI) was 623ms. Trails with different number of tones were randomly presented. The tone sequence could be in an alternating pattern (RE conditions in the upper two rows) or was presented randomly without any patterns (IR conditions in the lower two rows). The last stimulus in a trial was always a tone in noise with the individual measured SNR in the pre-test. The target tone was randomly selected from /ā/ or /á/, and was presented in noise in the same way as in the pre-test ---100 ms after the onset of 1s-long white noise and the target tone lasted for 377ms. Participants judged the target tone by pressing one of two buttons. The target tone in the RE conditions was either consistent (RE_con_) or inconsistent (RE_inc_) with the alternating pattern in the preceding clean tones. Whereas the target tone in the IR conditions was either the same (IR_same_) or different (IR_diff_) from the tone immediately before.

We manipulated two parameters in this procedure to investigate how the top-down prediction interacted with the bottom-up sensory processing and influenced the perceptual sensitivity and bias. The first parameter was whether the clean tone sequence was presented in an order. The sequence could be a regular pattern (RE) in which two tones are presented in an alternating manner. Or the sequence could be constructed by randomly presenting the two tones (IR). The second parameter was whether the tone in noise was consistent with prediction of different time-scales. In the RE conditions, the last tone in noise could be consistent with the regularity (RE_con_) or inconsistent (RE_inc_). That is, whether the tone of test was consistent with the long-term prediction formed by the regularity of preceding tone sequences. Whereas, in the IR conditions, the tone in noise could be the same as (IR_same_) or different from the last clean tone (IR_diff_). That is, whether the target tone was a repetition that was consistent with the short-term immediate effect formed by the last clean tone in a random sequence. Therefore, a total of four conditions were included in this experiment. Thirty-two trials were included for each condition, yielding a total of 128 trials. The presentation order of trials was pseudorandom across all participants.

### Statistical analysis

When computing the measures to quantify responses, we took the tone /á/ as the target tone. Hit rate was calculated as a proportion of correct response on the /á/ tone, while the proportion of making /á/ responses to the /ā/ tone stimulus was defined as the false alarm rate. Following the Signal Detection Theory, the detection sensitivity (or discrimination ability) can be expressed by calculating the sensitivity index (d’) (Macmillan and Creelman, 2004).

Statistical analyses were performed using IBM SPSS Statistics version 17.0, GraphPad. Prism 5.02. The normal distribution of data was tested using the Kolmogorov-Smirnov tests. One-way analysis of variance (ANOVA) was performed with factors of group for demographic and clinical continuous variables and chi-squares (χ^2^) test for categorical values. A univariate analysis of covariance (ANCOVA) was used to assess the performance of the different subject groups in four conditions. Pearson correlation analyses were performed to determine the relationship between clinical variables and behavioral data within AVHs patients. We used stepwise multiple regression analysis with d’ scores as the dependent variable to investigate the impact of age, gender, age of onset, duration of illness, AHRS total scores and PANSS and its subscales. Data are presented as mean (SD). Differences at *p* < 0.05 were considered to be significant.

## Results

### 1. Demographic data

Table 1 shows the subjects’ demographic data and the clinical variables. ANOVA analyses showed a significant difference in age (*p* < 0.001) among three groups, but not in education, height, and weight (p > 0.05). The χ^2^ test showed no significant differences among three groups with regard to gender (χ^2^ = 0.68, *p* = 0.508). Further, P3 subscore, positive and general psychopathology subscores were significantly higher in AVHs patients than non-AVHs patients (all *p* < 0.01). Neither the age of onset, duration or the PANSS total score was significantly different between two patients’ groups.

**Table1.** Demographics of Schizophrenia patients with and without AVHs and Healthy Controls.

| | AVHs Patients<br>(n=32) | Non-AVHs Patients<br>(n=29) | Healthy Controls (n=30) | F or $\chi^2$ | p-value |
| --- | --- | --- | --- | --- | --- |
| **Gender (M/F)** | 14/18 | 12/17 | 9/21 | 0.68 | 0.508 |
| **Age (years)** | 27.28 (6.37) | 25.79 (5.94) | 21.67 (2.45) | 9.41 | <0.001 |
| **Education (years)** | 12.44 (2.47) | 12.83 (2.65) | 13.43 (0.77) | 1.69 | 0.19 |
| **Height (m)** | 1.68 (8.31) | 1.66 (9.08) | 1.67 (7.07) | 0.39 | 0.677 |
| **Weight(kg)** | 66.78 (15.43) | 62.38 (7.10) | 60.37 (8.93) | 2.66 | 0.076 |
| **Age of onset (years)** | 21.09 (6.66) | 19.69 (4.80) | N/A | 0.82 | 0.368 |
| **Duration (month)** | 74.05 (77.64) | 63.38 (52.53) | N/A | 0.39 | 0.536 |
| **PANSS total score** | 82.03 (8.87) | 78.55 (4.41) | N/A | 1.97 | 0.06 |
| **P3 subscore** | 4.75 (0.98) | 1.00 (0.00) | N/A | 21.56 | <0.001 |
| **P subscore** | 23.53 (3.82) | 14.69 (2.65) | N/A | 10.40 | <0.001 |
| **N subscore** | 19.78 (3.48) | 22.41 (1.43) | N/A | 3.93 | 0.001 |
| **G subscore** | 38.72 (4.09) | 41.45 (2.56) | N/A | 3.09 | 0.003 |
| **AVHs total score** | 25.25 (3.85) | N/A | N/A | N/A | N/A |

Notes: PANSS: Positive and Negative Syndrome Scale; P subscore: positive symptom subscore;

### 2. Performance on the speech tone recognition task

The sensitivity indices d’ scores were first subject to a two-way mixed ANOVA with groups (AVHs, non-AVHs, and Healthy Control) as a between-subject factor and conditions (RE_con_, RE_inc_, IR_same_, and IR_diff_) as a within-subject factor. The omnibus results showed that the main effect of the groups was significant (*F*_(2,352)_ = 5.089, *p* = 0.007). The main effect of conditions was also significant (*F*_(3,352)_= 8.953, *p* = 0.001). More importantly, the interaction was significant (*F*_(6,352)_= 5.315, *p* = 0.001).

To further explain the interaction, we compared tone recognition between conditions within each group to investigate how predictions of different time scales modulated perception in different populations. Sensitivity indices d’ were calculated for the four conditions in the AVHs group (Figure 2A), non-AVHs group (Figure 2B) and healthy control group (Figure 2C). In the AVHs group, as shown in Figure 2A, the d’ scores in the RE_con_ condition was significantly higher than that in the RE_inc_ condition (*t*_(1,62)_ = 7.45; *p* < 0.001). However, the d’ was not different between the IR_same_ and IR_diff_ conditions, suggesting the tone recognition was not influenced by the immediate memory effect. This observation was consistent with our hypothesis suggesting that the long-term prediction biases perception in patients with AVHs.

**Figure 2.**
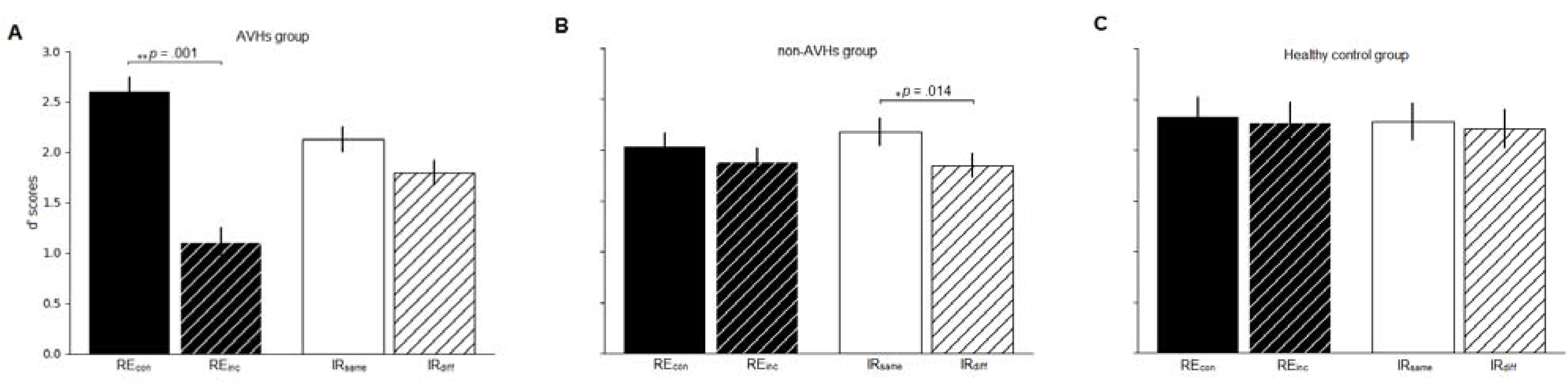
Tone identification results of three groups. A) Identification sensitivity index, d’ results in four conditions from the group of patients with auditory verbal hallucinations. B) Results from the group of patients without auditory verbal hallucinations. C) Results from the goup of health participants. Error bars indicate standard errors. **significant at the level of p < 0.01; *significant at the level of p < 0.05.

In non-AVHs patients, the d’ scores in the conditions with regular sequences, no significant difference was found between the Re_con_ and RE_inc_ conditions. Whereas in the IR_same_ condition was significantly higher than that in the IR_diff_ condition (*t*_(1,56)_ = 2.62; *p* = 0.014). These results directly contrast with the results in the AVHs group, suggesting that the perceptual judgment in the non-AVH group was more influenced by the immediate memory effect.

In the healthy control group, neither the difference between RE_con_ and RE_inc_ nor the difference between IR_same_ and IR_diff_ was significant, suggesting that healthy controls made perceptual judgment without influences from long-term predictions or short-term memory effect.

### 3. The relationship between clinical variables and behavior data of the speech tone recognition task

As shown in Figure 3A, a significantly positive correlation was found between AHRS total scores and d’ scores of RE_con_ condition in AVHs patients group (*r* = 0.576, *p* < 0.001). Further stepwise multiple regression analysis identified the AHRS total scores as a significant predictor for d’ scores in the RE_con_ condition (beta = 0.113, *t* = 3.607, *p* = 0.001) in AVHs patients. Whereas the other variables showed no effects (*p* > 0.05). The difference in d’ scores between RE_con_ and RE_inc_ was computed within AVHs groups to indicate the total influence of regularity on perceptual judgment. The following correlation analyses showed a significantly positive correlation between the d’ difference and the AHRS total scores (Figure 3B; *r* = 0.771, *p* < 0.001). Moreover, the PANSS P3 subscores were also significantly positively correlated with the d’ difference (Figure 3C; *r* = 0.453, *p* = 0.009). No significant correlation between the d’ difference and PANSS total, positive, negative and general psychopathology subscale scores (all *p* > 0.05). No significant correlation was found in non-AVHs patients. These results suggest that the degree of severity in AVH relates to the impacts of memory on perception.

**Figure 3.**
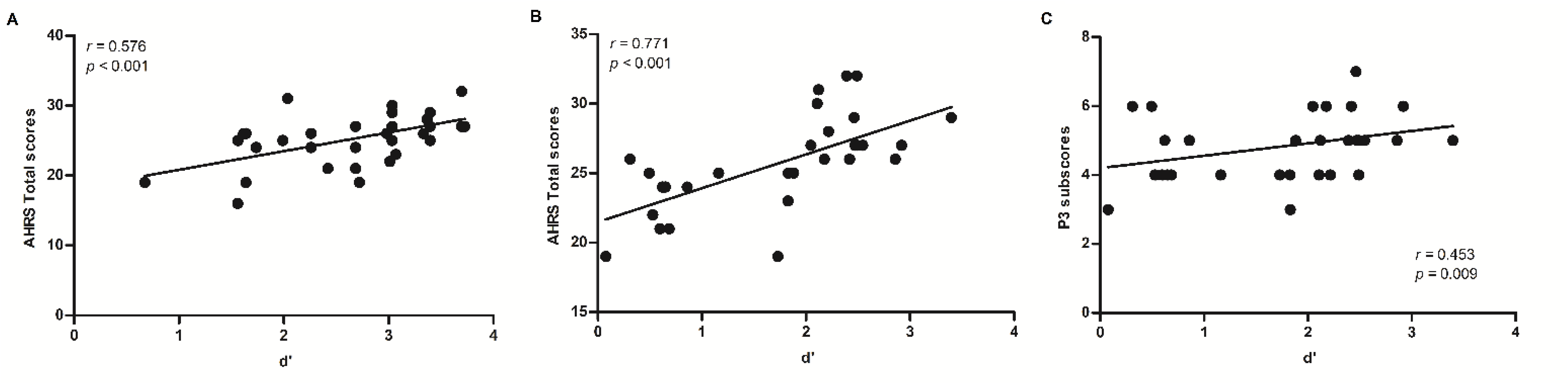
Results of correlations between tone identification and indices of symptom severity. A) The correlation between AHRS total scores and d’ in the RE_con_ condition in AVHs patients. B) The correlation between AHRS total scores and d’ difference between RE_con_ and RE_inc_ condition in AVHs patients. C) The correlation between P3 subscorses and d’ difference between RE_con_ and RE_inc_ condition in AVHs patients. AHRS, auditory hallucinations rating scale.

## Discussion

We investigated the effects of top-down predictions on perceptual processing of tones in schizophrenia patients with and without AVHs. We found that AVHs patients identified tones better when predictions met external stimuli, whereas performance was deleterious when prediction and stimuli were inconsistent. Moreover, the modulation effects were prominent for the predictions that were derived from long-term regularities in AVHs patients. In contrast, patients without AVHs showed the effects of short-term predictions from immediate repetitions. These consistent results collaboratively revealed the featural and temporal characteristics of weighting function in source monitoring of AVHs. Malfunctioned source monitoring in AVHs heavily weights predictions to form incorrect perception. The weighting function in source monitoring can extend to tonal features and predictions at multiple timescales differentially modulate perception in different clinical populations.

In this study, we extended the investigation of AVHs to basic featural level of tones. By manipulating the prediction as a function of expectancies in a trial-to-trial probabilistic fashion, we found that in the RE_con_ condition where the target tone was consistent with regularity of preceding tone sequence, AVHs patients had higher hit rate and lower false alarm rate, so that the sensitivity indices d’ became higher (Figure 2A). Moreover, the severity of AVHs positively correlated with the modulation effects of prediction (Figure 3A). In the RE_inc_ condition where the target tone violated the regularity, the AVHs patients produced more false positives, so that the sensitivity indices d’ became lower. The observations of more false positives are consistent with that verbal imagery and expectation cause more false positives of hearing speech in white noise in hallucination-prone participants (Moseley *et al.*, 2016, Vercammen and Aleman, 2008). The ‘apparent’ benefit of prediction in the RE_con_ condition and deleterious effect of prediction in the RE_inc_ condition are actually the results of confusing internal and external sources. The AVHs patients weighted more on the internal prediction but cannot correctly perform the sensory analysis which is the relevant processing in the tone identification task. These results are consistent with our hypothesis that tonal representations induced by prediction occupy neuronal resources of auditory cortex, making it less responsive to external stimulation. That is, AVHs may be ‘parasitic’ memories due to disrupted language production processes that spontaneously and erroneously activate language based memory (Hoffman, 1986, Li *et al.*, 2020, Ma and Tian, 2019, Tian and Poeppel, 2012). Moreover, these results suggest that the incorrect weighting in the source monitoring can extend to basic sound features of frequency.

Interestingly, a double dissociation was found between AVHs and non-AVHs groups regarding the effects of long-term and short-term predictions (Figure 2A.2B). Theoretically, in neural circuits, short-term and long-term plasticity of synaptic efficacy in sensory and motor neurons support learning and memory (Hasson *et al.*, 2008). Prior information can reshape synapses over different timescales by changing levels of activation, excitability and potentiation over milliseconds and minutes (Mongillo *et al.*, 2008, Perrett *et al.*, 2009). Thus, the synaptic plasticity can be a neural mechanism for continuously integrating prior information into processing of incoming information. Cognitively, the levels of information (e.g. phonemes, syllables, words, sentences) can be bases for forming predictions at multiple timescales that influence processing of incoming information along the speech hierarchy (Hasson *et al.*, 2010). The neural and cognitive foundations enable predictions form in different timescales and influence perception.

The distinct modulation effects of predictions at multiple timescales suggest separate mechanisms in different clinical populations. Contrasting with AVHs patients who showed long-term prediction effects, the non-AVHs patients tend to judge the target tone the same as the one immediately before. These results suggest that non-AVHs patients are prone to the influences of short-term memory. Schizophrenia patients without AVHs may have a greater response bias, rather than perceptual sensitivity deficits. These results are consistent with the involvement of externalizing biases in schizophrenia (Bentall and Slade, 1985, Varese *et al.*, 2012). Patients develop an external attribution bias as an explanation for the confusing abnormal perceptual/cognitive experiences of psychosis. This process may be related to the conscious evaluation of the external stimuli, which is presumably caused by lacks of effective connectivity between superior temporal gyrus(STG) and anterior cingulate cortex(ACC) – a critical component of the ‘core control network’.

In summary, we found that schizophrenia patients with AVHs weighted predictions more over sensory processing and altered the recognition of tones. Moreover, patients with and without AVHs showed distinct influences of predictions at different timescales. Our results support a Bayesian cognitive account that malfunctioned source monitoring mediate AVHs.

## Acknowledgments

We thank Jiaqiu Sun for his help on programing computer scripts, Yan Chen, Xinyu Fang and Dandan Wang for their help with recruitment and preprocessing of the data.

## Funding

This work was supported by the National Natural Science Foundation of China 31871131, Major Program of Science and Technology Commission of Shanghai Municipality (STCSM) 17JC1404104, Program of Introducing Talents of Discipline to Universities, Base B16018 to XT, and East China Normal University (ECNU) Academic Innovation Promotion Program for Excellent Doctoral Students (YBNLTS2019-026) to FY.

## Conflicts of Interest

None

## Data availability

The data that support the findings of this study are available from the corresponding authors on reasonable request.

## Author contributions

FY designed the experiment, collected and analyzed data, and drafted and edited the manuscript. HZ edited the experiment Python script and the methods. CZ, LY and WL, as psychiatrists, helped recruit schizophrenia patients. XT designed the experiment, interpreted the data, edited the manuscript, and provided critical revisions. All authors approved the final version.

## Ethical standards

The authors assert that all procedures contributing to this work comply with the ethical standards of the relevant national and institutional committees on human experimentation and with the Helsinki Declaration of 1975, as revised in 2008.

## Notes

### Competing Interest Statement

The authors have declared no competing interest.

